# The structural basis for the selective antagonism of soluble TNF-alpha by variable new antigen receptors

**DOI:** 10.1101/2024.07.30.605874

**Authors:** Obinna C. Ubah, Eric W. Lake, Stella Priyanka, Ke Shi, Nicholas H. Moeller, Andrew J. Porter, Hideki Aihara, Aaron M. LeBeau, Caroline J. Barelle

**Affiliations:** Elasmogen Ltd, Liberty Building Foresterhill Road, Aberdeen AB25 2ZP, UK; Department of Pathology and Laboratory Medicine, University of Wisconsin School of Medicine and Public Health, Madison, WI 53705, USA; Department of Biochemistry, Molecular Biology, and Biophysics, University of Minnesota, Minneapolis, MN 55455, USA; School of Medical Sciences, Scottish Biologics Facility, University of Aberdeen, Foresterhill, Aberdeen AB25 2ZP; Department of Radiology, University of Wisconsin School of Medicine and Public Health, Madison, WI 53705, USA; Carbone Cancer Center, University of Wisconsin-Madison, Madison, WI 53705, USA

## Abstract

The pro-inflammatory cytokine tumor necrosis factor-alpha (TNF)-α is synthesized as transmembrane TNF-α (tmTNF-α) where proteolytic processing releases soluble TNF-α (sTNF-α). tmTNF-α can act as either a ligand by activating TNF receptors, or a receptor that transmits outside-to-inside signals (reverse signalling) after binding to native receptors. All TNF-α therapies bind tmTNF-α and induce reverse signalling which can result in immunosuppression leading to infection. We present crystal structures of two anti-TNF-α Variable New Antigen Receptors (VNAR) in complex with sTNF-α *via* two distinct epitopes. The VNAR-D1 recognized an epitope that selectively engaged sTNF-α while VNAR-C4 bound an epitope that overlapped with other biologic therapies. In activated CD4^+^ T cells, our VNARs did not bind tmTNF-α in contrast to commercially available therapies that demonstrated induction of reverse signalling. Our findings suggest that neutralisation through a unique mechanism may lead to anti-TNF-α agents with an improved safety profile that will benefit high-risk patients.

## Introduction

Tumor necrosis factor-alpha (TNF-α) is a clinically relevant cytokine that regulates biological activities such as cell growth, carcinogenesis, inflammation, viral replication, immune surveillance, and autoimmunity.^1^ As a proinflammatory cytokine, TNF-α is a principal mediator of autoimmunity and inflammatory diseases, including Crohn’s disease (CD), ulcerative colitis (UC), and rheumatoid arthritis (RA). It exerts its function by binding to two structurally distinct membrane receptors, TNFR1 (also known as TNFRSF1A, CD120a) and TNFR2 (TNFRSF1B, CD120b). TNFR1-mediated signalling is predominantly associated with inflammation and cell death, whereas TNFR2 mainly regulates T-cell migration and tissue regeneration.^2,3^ Blocking the interaction of TNF-α with its two receptors has been exploited as a therapeutic mechanism for treating inflammatory and autoimmune diseases.^3–5^ Soluble TNF-α (sTNF-α) is generated through the proteolytic cleavage of its precursor form, transmembrane TNF-α (tmTNF-α), by the enzyme TNF-α converting enzyme (TACE or ADAM-17). Growing evidence also suggests that tmTNF-α is involved in the regulation of a local inflammatory response. tmTNF-α is expressed on most cell types including activated macrophages and lymphocytes, and it functions through cell-to-cell contact, contrary to sTNF-α which acts at sites distal from TNF-α producing cells.^6–8^ tmTNF-α has mechanistic duplexity as a ligand that binds to TNFR1/TNFR2 and as a receptor that conducts outside-to-inside (reverse) signalling on tmTNF-α bearing cells. It is therefore believed that tmTNF-α plays a critical role in local inflammation.^9–13^

Anti-TNF-α biologics have been successfully implemented in the clinic for the treatment of chronic inflammatory diseases such as RA, UC, CD, psoriatic arthritis (PsA), psoriasis (PsO), and ankylosing spondylitis (AS). However, these clinical successes have been accompanied by a two-to-four-fold increased risk of granulomatous infectious diseases, including active tuberculosis (TB), reactivation of latent TB, invasive fungal infections, and bacterial and viral infections by a range of opportunistic pathogens.^11,14,15^ There also appears to be differences in the clinical efficacy of the anti-TNF-α drug class for certain inflammatory disease indications. For instance, anti-TNF-α antibodies Infliximab, Adalimumab and Etanercept are equally effective in RA, but only Infliximab and Adalimumab are effective in inflammatory bowel diseases. Therefore, besides the common sTNF-α neutralisation, additional functional features such as biology resulting from interaction with tmTNF-α or Fcγ receptor-expressing cells can result in degradation and inflammation of tissue and can contribute to the overall clinical and adverse events documented for each anti-TNF-α biologic (**Figure 1**). Some of these adverse events may be explained by the binding of anti-TNF-α biologics to tmTNF-α, and the triggering of “reverse signalling” that induces cell-cycle arrest, suppression of T-cell proliferation and apoptosis of tmTNF-α bearing cells.^16–22^ It is likely that the induction of tmTNF-α mediated reverse signalling by itself does not explain all reported adverse events with anti-TNF-α agents, but manipulation of this characteristic of anti-TNF therapies provides a route to rationally design safer and more broadly effective anti-TNF-α agents with minimal or abrogated tmTNF-α interaction. Such an approach may reduce overall side-effects from TNF therapies and may also provide, for the first time, access to anti-TNF drugs for patient groups that are currently unable to benefit from this drug class (e.g. patients with latent bacterial or viral infections, patients at risk of malignancies).

**Figure 1.**
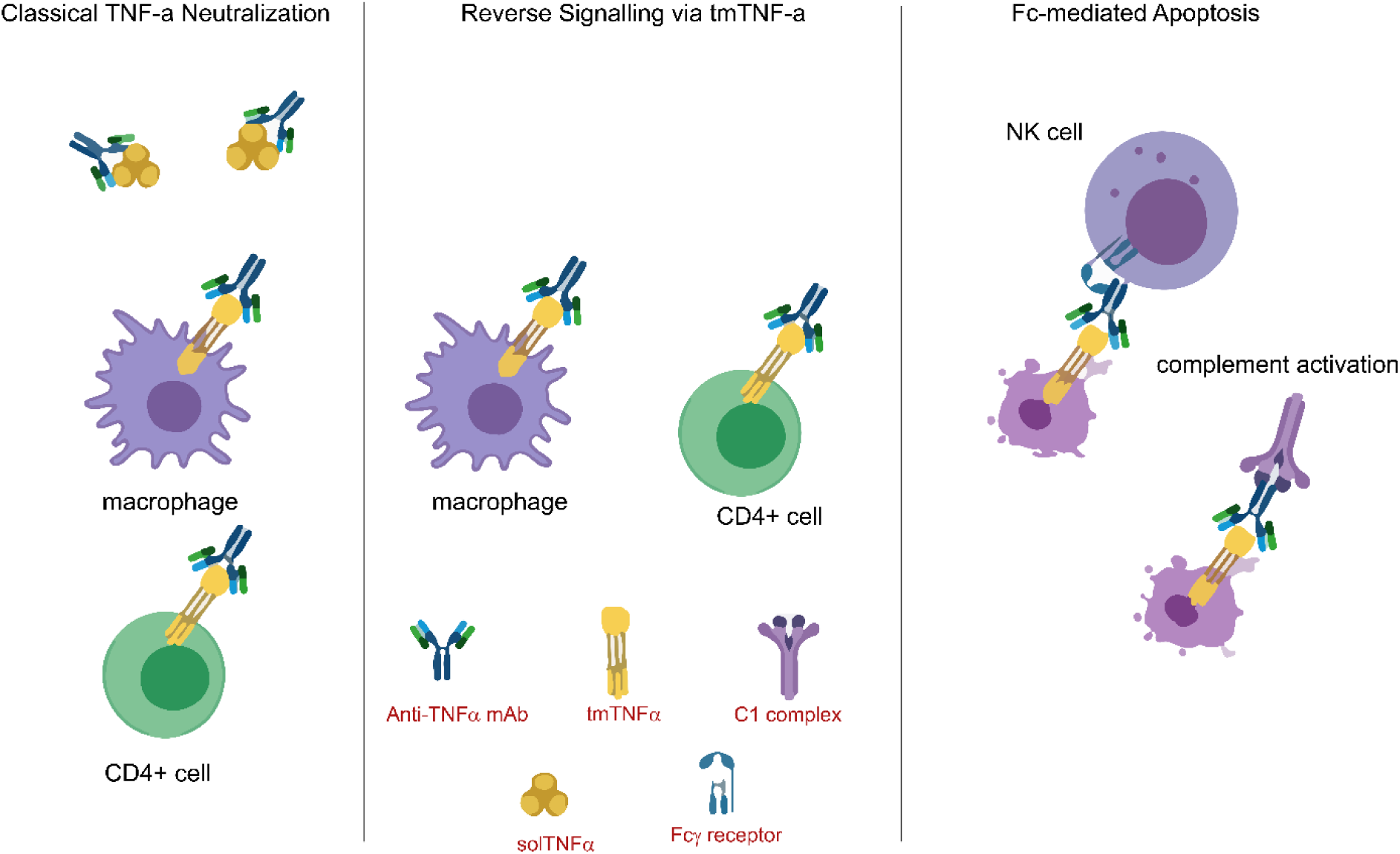
Anti-inflammation mechanisms utilised by anti-TNF-α agents. Broadly grouped into the **classical** neutralisation of soluble TNF-α (sTNF-α) and transmembrane TNF-α (tmTNF-α), Reverse signalling mediated by the neutralisation of membrane-bound TNF-α (tmTNF-α), and Fc-mediated apoptosis which is only triggered by those anti-TNF-α agents capable of binding both Fcγ receptors (FcγRs) and complements.

Variable New Antigen Receptors (VNARs) are small (∼11 kDa), naturally occurring independent binding domains isolated from the adaptive immune repertoire of sharks. One salient characteristic of VNARs is their unique targeting properties that allow the VNAR paratope to access and bind epitopes inaccessible to conventional biologics^23–29^. We have previously reported the isolation of high affinity, super-potent, clinically applicable anti-TNF-α VNARs (D1 and C4) that possess low or no immunogenicity in murine models.^25,30,31^ Here, we present co-crystallisation data of the anti-TNF-α VNARs complexed with TNF-α that provides evidence of a VNAR engaging an epitope that is novel among the currently published landscape of existing anti-TNF-α biologics. This recessed epitope on sTNF-α recognised by VNAR-D1 may be the ultimate discriminator between sTNF-α and tmTNF-α, consequently allowing for rational design of safer and broadly efficacious anti-TNF-α therapies.

## Materials & Methods

### Expression and purification of soluble TNF-α and VNARs

Synthetic genes encoding the soluble fragment of human TNF-α (residues 77-233 of the membrane bound form, uniprot ID PO1375), and the VNARs D1 and C4 with N-terminal, 3C protease-cleavable hexa-histidine tag were synthesised at GeneArt Gene Synthesis Thermo Fisher Scientific. *E. coli* BL21 Gold pLysS cells were transformed with expression vectors encoding human sTNF-α, VNAR D1 and VNAR C4 sequences with an N-terminal 3C protease-cleavable hexa-histidine tag. Positive transformants were selected on LB-agar containing appropriate antibiotic, and expression induced in LB medium using 0.5 mM isopropyl β-D-1-thiogalactopyranoside (IPTG). Harvested cells were resuspended in PBS supplemented with Complete protease inhibitors and 5 mM β-mercaptoethanol and lysed by three passes through an EmulsiFlex C5 homogenizer. Cleared lysates were mixed with Ni-NTA agarose beads equilibrated in PBS, and bound protein was eluted with 500 mM imidazole (prepared in 20 mM Tris HCl, pH 8.0, 150 mM NaCl, 5 mM β-mercaptoethanol (Buffer A). Eluted protein was cleaved with 3C protease-GST tag and dialysed overnight in 20 mM Tris HCl, pH 8.0, 150 mM NaCl, 5 mM β-mercaptoethanol buffer. Uncleaved protein, cleaved hexa-histidine tag and 3C protease-GST tag were removed by passing the dialysed sample through a Glutathione agarose and a Ni-NTA column. The untagged protein (sTNF-α, VNAR D1 and VNAR C4) flowthrough were further purified using cation exchange and/or size exclusion chromatography and dialysed into PBS. The VNAR Quad-X® constructs were designed, gene synthesised, expressed, and purified as previously described.^25,31^ VNAR Quad-X constructs are approximately 105 kDa fusion molecules designed with two varying lengths of glycine-serine (GlySer) linkers, a shorter 2 – 3 (Gly_4_Ser) fusing the first anti-TNF-α VNAR-D1 or C4 to the hinge of a wild-type human IgG1 Fc region, and a 4 – 5 (Gly_4_Ser) linker fused to the C-terminal of the human IgG1 Fc CH-3 domain linking the second anti-TNF-α VNAR-D1 or C4. NB: Co-crystallisation of the anti-TNF-α VNARs with soluble TNF-α was conducted using monomer VNAR-D1 and VNAR-C4.

### *In vitro* functional characterisation of the mono-paratopic Quad-X constructs

The TNF-α sensitive mouse fibrosarcoma cell line (L929 cells) were grown to 90% confluence, seeded onto 96-well flat bottom microtiter cell culture plate at 5,000 cells per well, and incubated for 48 h. Anti-TNF-α VNAR Quad-X were added to actinomycin-D and sTNF-α treated L929 cells. Cytotoxicity or cell survival was determined by adding tetrazolium salt (WST-1) cell proliferation reagent, and absorbance read at 450 nm. Octet R8 biolayer interferometry (BLI) was used to determine binding kinetics of the mono-paratopic bivalent VNAR Quad-X to sTNF-α. Anti-TNF-α VNAR Quad-X prepared in 1% BSA in PBS + 0.01% Tween 20, pH 7.4 (assay buffer) were loaded on assay buffer-equilibrated anti-human IgG Fc Capture (AHC2) biosensors at a final concentration of 10 nM for 10 min, followed by 30 min association step with serial dilutions of sTNF-α prepared in assay buffer and a 30 min dissociation step. Data were analysed using the Octet® BLI Analysis Studio software 12.2., and global 1:1 fit model. Binding kinetics were also measured using biotin labelled sTNF-α prepared in assay buffer (1% BSA in PBS + 0.01% Tween 20, pH 7.4) at a final concentration of 10 nM. Biotinylated sTNF-α was loaded on assay buffer-equilibrated High Precision Streptavidin (SAX) biosensors for 10 min followed by 30 min binding association of serially diluted anti-TNF-α VNARs and a final 30 min binding dissociation step.

### sTNF-α – anti-TNF VNARs complex assembly & size exclusion chromatography (SEC) of complexes of TNF-α with VNARs

Complex assembly was first tested at microgram scale. Successful assembly protocols were scaled up to milligram amounts. At small scale tests, 100 µg sTNF-α (5.7 nmol) was mixed with a 2-fold molar excess (11.4 nmol) of VNAR in 100 µL Buffer A. sTNF-α-VNAR mixtures were incubated for 30 min on ice and centrifuged for 10 min, 21,000 xg, at 4 °C. Supernatants were analysed by SEC using Superdex 200 10/300 column, and SEC fractions were analysed by SDS PAGE. Fractions containing target complexes were pooled, concentrated, and used for crystallisation screens.

### Crystallisation and X-ray structures determination for VNAR-D1 and sTNF-α complex

Crystallisation conditions for sTNF-α-VNAR-D1 were screened following the sitting drop vapour diffusion protocol.^32^ Drops were monitored regularly using an optical microscope. Initial hits were refined in 48-well plates using hanging drop setups with 500 µL reservoir solution and 1 – 6 µL drops with streak seeding. The best crystals grew after several rounds of seeding in a condition containing 19 % PEG 8000, 0.2 M ammonium sulphate and 0.1 M MES pH 6.0 at 18 °C. Crystals were harvested with nylon loops, transferred to a cryoprotectant (reservoir solution supplemented with 10 – 20 % glycerol, ethylene glycol, propylene glycol or PEG 400) and flash-frozen in liquid nitrogen. X-ray diffraction data was collected at beamline P14, German Electron Synchrotron DESY, Hamburg, and a complete dataset was collected at 3.31 Å resolution (Table S1). Diffraction images were processed using XDS.^33^ Phases were obtained by molecular replacement with PHASER using the TNF trimer from Protein Data Bank (PDB) entry 1TNF^34^. The model was completed with COOT (Emsley, Cowtan 2004) and refined using REFMAC5.^35,36^ Protein-protein interfaces were analysed using PISA. PyMOL and COOT were used for refinement of the crystal structure and comparison of the sTNF-α-VNAR-D1 structure to other known soluble huTNF-α/anti-TNF-α complexes. The VNAR-D1: sTNF-α crystallisation and X-ray structural determination was carried out at moloX GmbH, Berlin Germany.

### Crystallisation and X-ray structures determination for VNAR-C4 and sTNF-α complex

Purified sTNF-α and VNAR-C4 were mixed at ∼1:1 molar ratio and the complex formed was isolated using a Superdex200 size-exclusion chromatography (SEC) column operating with 10 mM Tris-HCl (pH 7.4) and 150 mM NaCl. The isolated complex was concentrated by ultrafiltration to ∼10 mg/ml and subjected to crystallization. The sTNF-α-VNAR-C4 complex was crystallized by the sitting drop vapor diffusion method, using a reservoir solution containing 0.1 M MES, pH 6, 0.1M magnesium chloride, 8% w/v polyethylene glycol 6000. The crystals grew to full size in three days. The crystals were cryo-protected by briefly soaking in the reservoir solution supplemented with 25% ethylene glycol and flash-cooled by plunging into liquid nitrogen. X-ray diffraction data were collected at the Advanced Photon Source NE-CAT beamline 24-ID-C. All X-ray diffraction data were processed using XDS version 20210205. Molecular replacement calculations were performed using PHASER version 2.8.^34^ Iterative manual model building and refinement were done using COOT version 7766 and PHENIX version v.1.19.2-4158, respectively.^37,38^ The summary of data collection and model refinement statistics is shown in Table S2. The VNAR-C4:sol huTNF-α crystallisation and X-ray structural determination was carried out at the Department of Biochemistry, Molecular Biology, and Biophysics, University of Minnesota, Minneapolis. The coordinates and structure factors will be deposited in the Protein Data Bank on May 31, 2024.

### Transmembrane TNF-α binding and reverse signalling in phytohemagglutinin (PHA) activated CD4+ primary human T cells

Healthy human CD4^+^ T cells (BioIVT, lot no. HumanCD4-0111394) were cultured in supplemented RPMI at density of 0.2 x 10^6^ cells/mL in 96-well flat bottom cell culture plates, 37 °C, 5 % CO_2_. CD4^+^ T cells were stimulated with 0.1 µg/mL PHA for 12 h. PHA activated cells were washed with acidified buffer (50 mM glycine-HCl containing 150 mM NaCl, pH 3.0) for 3 min at 4 °C to remove traces of sTNF-α. Acidified buffer was removed by washing the cells three times with PBS containing 1 % BSA. PHA activated cell were treated with anti-TNF-α VNARs at different concentrations and incubation timepoints. Cells were stained for E-selectin using anti-human CD62E-FITC conjugated antibody and fixed with 0.5 % paraformaldehyde (PFA) for 15 min, 4 °C, and fluorescence intensity measured using CLARIOstar® *plus* microplate reader and Attune™ NxT flow cytometer. IFN-γ levels was measured in anti-TNF-α treated activated CD4^+^ T cells supernatant using the human IFN-γ DuoSet Sandwich ELISA kit (R&D systems, Cat no. DY285B-05).

## Results

### VNAR D1 recognizes a novel epitope only accessible on soluble TNF-α

VNARs C4 and D1 were separately complexed with sTNF-α (aa residues 77-233) and purified by size-exclusion chromatography. Both complexes were subjected to crystallization and yielded crystals. The structures were determined by molecular replacement phasing and refined to 1.99 and 3.31Å resolution for C4 (**Figure 2A**) and D1 (**Figure 2B)** respectively. In each structure, VNAR monomers were bound to each protomer of the TNF-α trimer in a 1:1 stoichiometry (**Figures 2A & B**). Analysis of the crystal structures revealed that VNAR-C4 binds to an epitope located near the C-terminal end of the TNF-α trimer in a similar location to other antibody therapeutics such as Adalimumab and Infliximab (**Figures 2 & S1A**). When compared with the contact binding surface of Adalimumab, which spans across multiple protomers and covers an area of 2540 Å^2^, the binding surface of VNAR-C4 covers only 822 Å^2^ on the surface of one protomer (**Figure 2C**).^39^ This highlights the ability of VNARs to achieve comparable binding affinities to conventional full-length immunoglobulins, but across a much smaller molecular footprint. Interestingly, VNAR-D1 binds a different and potentially unique epitope, not represented by any known antibodies **(Figure S1**). This epitope is in the groove formed between two adjacent protomers and is enveloped by the N-terminal tail of soluble TNF-α (**Figure 2B**). The binding surface of VNAR-D1 is also relatively small covering only 968 Å^2^ across the sTNF-α cleft, with high-affinity neutralisation facilitated by a finger-like binding loop that reaches into grooves on the target proteins.

**Figure 2.**
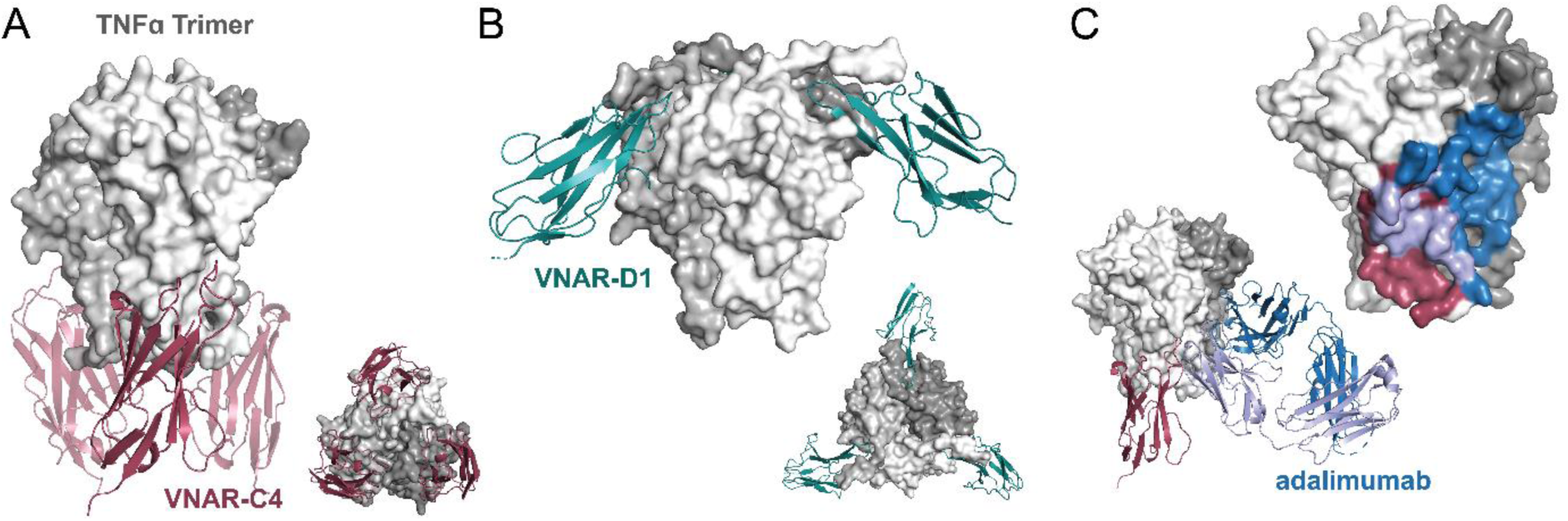
X-ray structures of VNARs expose novel TNF-α epitopes. The panels A and B show the crystal structure of soluble TNF-α trimer (grayscale) bound by multiple monomers of VNAR-C4 (A, red) and VNAR-D1 (B, green). Insets show a second view to highlight the 1:1 stoichiometry of TNF-α trimers to VNARs. C) The figure shows a grayscale surface representation of the TNF-α trimer with the epitopes for VNAR-D1 (red) and adalimumab (blue). Epitope overlap is colored light blue. Inset shows cartoon representations of each biologic bound to TNF-α as in Figure S1.

The structures of VNARs C4 and D1 each represent a facet of shark-derived therapeutics that apply to multiple disease contexts. Our studies with VNARs have so far discovered that VNARs either 1) possess similar mechanisms of action to existing biologics but achieved with a much smaller binding footprint, or 2) reveal/expose unique mechanisms of actions that make VNARs viable therapeutic candidates for patient groups that may not respond well to or are put at risk by existing therapeutic strategies.^24^ Here, the epitope of VNAR-C4 and its mechanism of action overlaps with that of the TNF receptors 1 and 2 (TNFR1 & TNFR2) directly competing with receptor binding (**Figure 3A**). By transposing the C4: TNF-α complex onto the structure of TNFR2 bound to TNF-α, shows C4 completely obscures the interaction of TNFR2 with TNF-α. This receptor-blocking action of C4 also holds true when the C4: TNF-α complex is transposed onto the structure of TNFR1 bound to TNF-α, with the caveat that the TNFR1: TNF-α structure used in this comparison was taken from a complex that was solved with an asymmetric form of the TNF-α trimer.

**Figure 3.**
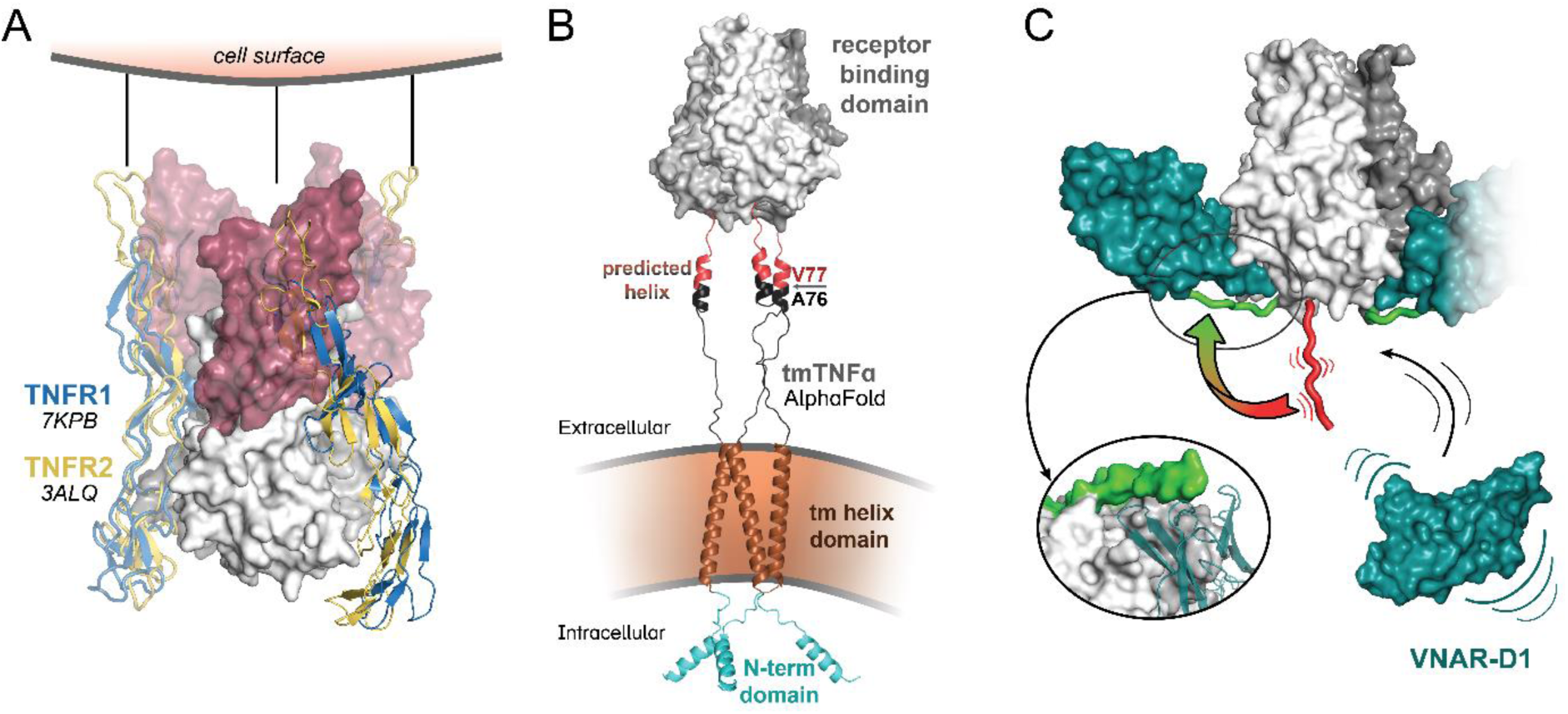
Novel VNAR mechanism of action against soluble TNF-α. A) Overlaid crystal structures of TNFR1 (blue) and TNFR2 (yellow) shown as cartoons bound to a surface representation of the TNF-α trimer (grayscale) with an illustration indicating the receptor position relative to the cell surface. A surface representation of VNAR-C4 has been superimposed onto the structures showing the overlap with the TNFRs. B) The figure shows a model of the tmTNF-α trimer tethered to a representation of the cell membrane with individual domains of TNF-α colored as follows: intracellular N-terminal domains=cyan, transmembrane helices=brown, spanning residues=black, RSSSRTPS-motif=red, receptor binding domain=-gray. C) The panel shows a schematic of the steps of free solTNF-α (grayscale) being bound by free VNAR-D1 monomers with motion lines indicating the conformational flexibility of the RSSSRTPS-motif (red). A red/green gradient arrow indicates that conformational change of the RSSSRTPS-motif as it envelopes VNAR-D1 in the bound state with an arrow to an inset showing the full transition.

VNAR-D1 also overlaps with the TNFR1 and TNFR2 binding site leading to receptor blocking activity that is synonymous with that of VNAR-C4 (**Figure S2A**). However, the VNAR-D1 epitope suggests a unique mechanism of action (condition 2 above) that allows selective targeting of the soluble form of TNF-α. To understand how VNAR-D1 selectively binds sTNF-α, it is important to understand the process by which sTNF-α is generated from tmTNF-α.^40^ After translation, full-length monomeric TNF-α is trafficked to the cell surface where it quickly forms a trimer, even at low nanomolar concentrations. The structure of full-length tmTNF-α has never been solved but has been predicted by AlphaFold as monomer. ^41,42^ By duplicating this form and rearranging the transmembrane tether, a model of trimeric tmTNF-α was constructed to visualize how tmTNF-α might look on the cell surface. AlphaFold predicts that a helix resides in the span of the residues preceding the sTNF-α receptor binding domain (**Figure 3B**), and that this same helix is also predicted by the Chou and Fasman secondary structure prediction (CFSSP) server when the sequence was uploaded. Directly in the middle of the predicted helix lies the cleavage boundary (residues A76 and V77, **Figure 3B**, red/black) where ADAM-17 canonically cleaves tmTNF-α into sTNF-α, exposing residues 77-85 of the N-terminal tail (RSSSRTPS-motif). Once sTNF-α is released from its cell surface tether, it can freely move and reorientate, allowing the RSSSRTPS-motif to fold around VNAR-D1 when it binds its epitope (**Figure 3C**). In contrast, when sTNF-α is tethered to the transmembrane portion of TNF-α, the residues of the RSSSRTPS-motif are wound into a helix, blocking the conformational change necessary to envelop the VNAR-D1 binding site. The RSSSRTPS-motif is essential for the high affinity interaction of VNAR-D1 with sTNF-α, and this requirement is also evident in the crystal structures. The B-factor values show that the electron density of the RSSSTRPS-motif is well ordered when stabilized by VNAR-D1, and even though an identical sTNF-α construct (residues 77-233) was used for the VNAR-C4 complex, the RSSSRTPS-motif is unresolved in that structure. In fact, the RSSSRTPS-motif is not visible in any published structures of TNF-α, demonstrating that it can only be visualized when fully stabilized. This suggests a completely novel mechanism of action of TNF-α neutralization and one that may offer new therapeutic possibilities.

### Analysis of the VNAR CDR3 engagement with the “unique” epitopes on the surface of TNF-α

The presentation of binding loops on a VNAR allows them to pack tightly into their epitope, with their protruding CDRs reaching into structural features that would be difficult to access by larger antibodies with planar binding faces formed from two separate binding domains. Covering a total area of 970 Å^2^ on the TNF-α trimer, the VNAR-D1 epitope is unique among existing anti-TNF-α biologics, such as adalimumab (**Figure S1**). This is because the VNAR-D1 epitope is in the cleft of two protomers, with one protomer interface accounting for 634 Å^2^ and another accounting for the remaining 336 Å^2^. This interface is repeated in triplicate on the whole TNF-α trimer allowing up to three VNAR-D1 molecules to engage. The “fingertip” of the VNAR-D1 CDR3, which consists of residues Gly90, Leu91, and Ala92, reaches deep into the cleft formed between two protomers of the TNF-α trimer and is enveloped by the residues of the RSSSRTPS-motif (**Figure 4A).** The hydrophobic fingertip engages with hydrophobic residues in the cleft, primarily with Val89, Ala114, and Leu112 of the first protomer of TNF-α (**Figure 4A**, *white)* as well as Leu131 of the second protomer (**Figure 4B**, *yellow*). The sidechain carbon segments of Thr83 and Ser85 of the first protomer and Gln203 of the second protomer are also oriented towards the hydrophobic finger, helping to stabilize the packing of VNAR-D1 into the cleft **(Figure 4A & B**, *yellow* and *white*). Of note, Leu91 of VNAR-D1 contributes 1.57 kcal/mol of solvation free energy to this interaction, which accounts for approximately one order of magnitude of the overall affinity. The N-terminal Arg78 of TNF-α at the end of the RSSSRTPS-motif, donates two hydrogen bonds (h-bonds) to Thr31 and Ser32 of VNAR-D1 (**Figure 4A**) and its carbonyl receives an h-bond from Lys64 of VNAR-D1 (**Figure 4B**). A substantial interaction accounting for a total of -2.03 kcal/mol of solvation energy is the result of Arg107 of the first TNF-α protomer coordinating a h-bond network with the sidechains of Glu86, Glu93, and Asp95 of VNAR-D1. Also, on the CDR3 finger, Tyr89 receives a h-bond at its carbonyl group from Ser81 of the RSSRTPS-motif of protomer 1, and it also forms an h-bonds with Glu129 of the second protomer. In addition, and to reinforce the strength of this interaction, the VNAR-D1 CDR3 backbone amine of Gly90 donates an h-bond to Gln201 of the second TNF-α protomer. Outside of CDR3, Arg25 of VNAR-D1 can donate an h-bond to Thr165 (**Figure 4B**). The carbonyl of Ser162 of the second protomer receives multiple h-bonds from both the backbone amine and the sidechain of VNAR-D1 Ser27 (**Figure 4B**). VNAR-D1 His26, depending on its sidechain orientation, can donate a h-bond to Glu203 or receive an h-bond from Ser162 of the second TNF-α protomer (**Figure 4B**). In total, the majority of the VNAR-D1 interaction occurs with CDR3, but its small profile allows multiple interactions across its surface.

**Figure 4.**
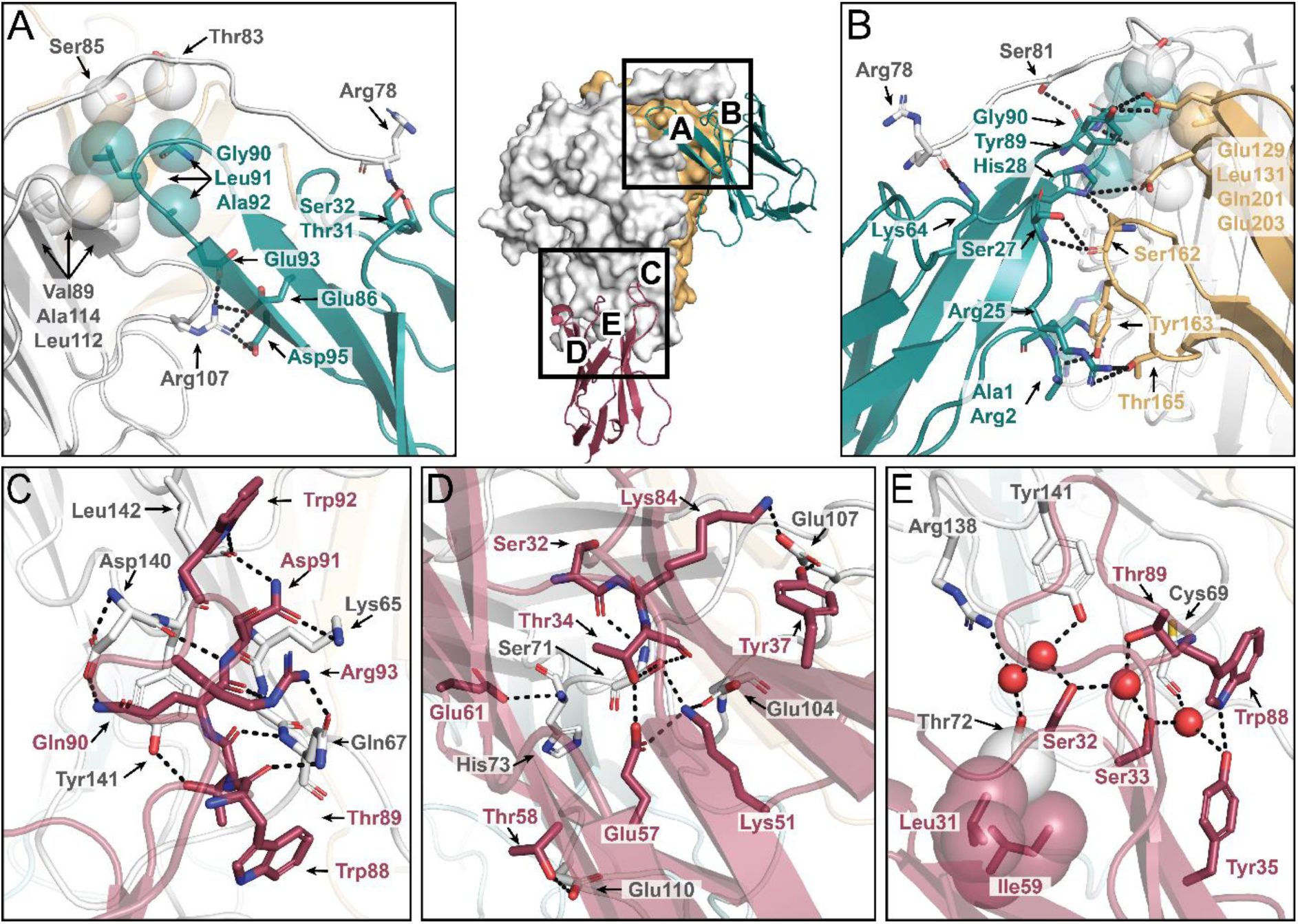
VNAR:TNF-α interactions are mediated primarily through the CDR3. The center of the figure shows an interaction map as a surface representation of the TNF-α trimer with protomer 1 colored white, protomer 2 colored yellow, and protomer 3 colored pale cyan. VNARs are represented as cartoon structures with VNAR-D1 colored green and VNAR-C4 colored red. Square boxes indicate the binding interfaces with letter labels corresponding to the approximate location of each of the zoomed in figures. A) Cartoon representation of a view of the VNAR-D1:TNF-α interface with sticks showing interacting residues with dashed lines representing hydrogen bonds and spheres representing hydrophobic interactions. Residue labels are colored to match their structures which are colored as in the central interaction map. B) A view of the VNAR-D1interaction surface rotated 180 degrees from panel A. Residue labels are colored to match their structure which is colored as in the central interaction map. C,D) A view of the VNAR-C4:TNF-α interaction with sticks representing interacting residues and dashed lines representing hydrogen bonds. Residue labels are colored to match their structures which are colored as in the central interaction map. E) A view of the VNAR-C4:TNF-α interaction shown as in panels C and D with added transparent spheres over residues with hydrophobic interactions. Water molecules are shown as solid red spheres with dashed lines indicating hydrogen bonds with residues. Residue labels are colored to match their structures which are colored as in the central interaction map.

In contrast, VNAR-C4 binds an epitope that overlaps with most clinically approved biologics targeting TNF-α (**Figure S1**). The epitope covers a single protomer that is accessible on all three protomers allowing VNAR-C4 to bind in triplicate. The surface area of the interaction covers 766 Å^2^. Here again, the interaction interface of VNAR-C4 highlights the unique finger-like nature of the CDR3 loop, with residues 88 – 93 engaging with a single protomer of TNF-α (**Figure 4C**). The carbonyl of Trp88 receives an h-bond from the sidechain of TNF-α Gln67; TNF-α Gln67 receives another h-bond from VNAR-C4 Arg93, whose sidechain is stabilized by an internal h-bond with the carbonyl of Gln90. Gln67 of TNF-α also donates an h-bond to the carbonyl of VNAR-C4 Thr89 forming a backbone-backbone interaction common among VNARs.^24^ This same backbone-backbone interaction also takes place between the amine of VNAR Asp91 and the carbonyl of TNF-α Asp140. The sidechain of VNAR-C4 Asp91 receives an h-bond from TNF-α Lys65, and, together with VNAR-C4 Trp92, donates an h-bond to the backbone carbonyl of TNF-α Leu142 (**Figure 4C**). And lastly, the sidechain of VNAR-C4 Gln90 interacts with the sidechain of TNF-α Asp140, which in turn is stabilized by an intra-residue h-bond with its backbone amine.

While CDR3 residues are the main drivers of binding, additional residues, outside this loop region, support the final interaction. VNAR-C4 residues Tyr37 and Lys84 each form an h-bond with the sidechain of TNF-α Glu107, and the sidechain of VNAR-C4 Glu61 accepts an h-bond from the backbone amine of TNF-α His73 (**Figure 4D**). VNAR-C4 Thr58 is the only residue in the paratope that interacts with a different protomer in the epitope: Glu110 of the third protomer of TNF-α. Lys51 of VNAR-C4 Lys51 donates two intermolecular h-bonds to the sidechains of TNF-α Ser71 and Glu10 respectively, but it also donates a stabilizing intramolecular h-bond to VNAR-C4 Glu57, forming a nearly perfect tetrahedral bonding geometry. A second but similar h-bonding geometry occurs from the side chain of TNF-α Ser71 which accepts two h-bonds from VNAR-C4 Thr34 and Lys51, while donating its hydrogen to the carbonyl of VNAR-C4 Thr34. Two additional interactions further stabilize this dual-tetrahedral h-bond network with the backbone amine of TNF-α Ser71 donating an h-bond to the carbonyl of VNAR-C4 Ser32, and VNAR-C4 Glu57 forms an intramolecular interaction with Thr34 (**Figure 4D**). Though the VNAR-C4:TNF-α interaction is primarily h-bond mediated, there is a subtle hydrophobic attraction between VNAR-C4 residues Leu31 and Ile59 with the methyl group of TNF-alpha Thr72 (**Figure 4E**). This epitope:paratope space also contains a water-mediated hydrogen bond network between VNAR-C4 residues Ser32, Ser33, Tyr35, Trp88, and Thr89 with TNF-α Cys69, Thr72, Arg138, and Tyr141 (**Figure 4E**). This water network is quite stable due to the favourable h-bond angles and the fact that the spatial location of the water molecules is conserved across all three interfaces in the crystal structure.

### Functional characterisation of the mono-paratopic tetravalent human Fc fusion VNAR constructs

The *in vitro* neutralising potency and TNF-α binding affinities of the monomeric VNAR D1 and C4 and a corresponding bi-paratopic tetravalent human Fc (huFc) fusion proteins (also known as Quad-X ELN22-108) consisting of 2x D1 and C4, have been previously reported.^25,31^ As the epitopes of VNAR D1 and C4 were confirmed through crystallography as being distinct, two mono-paratopic tetravalent Quad-X ELN22-129 (D1-huFc-D1) and ELN22-130 (C4-huFc-C4) were assessed for binding to sTNF-α. Given the loss of bi-paratopic binding, the mono-paratopic anti-TNF-α VNARs showed a predicted but acceptable loss in affinity and neutralising potency for sTNF-α compared to the parent bi-paratopic ELN22-108 molecule. We predicted this loss in affinity from previous work with the mono-paratopic and bi-paratopic dimers and trimers of the anti-TNF-α VNARs.^25^ The parent bi-paratopic ELN22-108 recorded affinity ranging between 9 and 74 pM depending on whether TNF-α is captured on the biosensor *via* biotin-streptavidin or presented in solution in the assay wells in 96-well plates **(Table 1**). ELN22-129 and ELN22-130 mono-paratopic constructs had affinities in the range of 10 – 230 pM, also depending on the antigen presentation approach. Affinities independently reported for Etanercept, Infliximab and Adalimumab vary from group to group, and are dependent on measuring instrument and assay method. However, affinities of these anti-TNF-α mAbs range between 23 pM – 1.15 nM.^19,39^ ELN22-108 had superior neutralising potency (EC_50_) of ∼ 3.9 pM compared to the 184 pM and 8.9 pM for mono-paratopic ELN22-129 and ELN22-130 respectively in the gold standard L929 cytotoxicity assay (**Figure 5**). There is a clear affinity and *in vitro* potency improvement achieved by targeting two distinct epitopes (ELN22-108) on TNF-α. The affinity obtained for the mono-paratopic VNARs in the Quad-X format are comparable with the values observed for the clinically approved anti-TNF-α biologics, Adalimumab, and Infliximab.^39,43^

**Figure 5.**
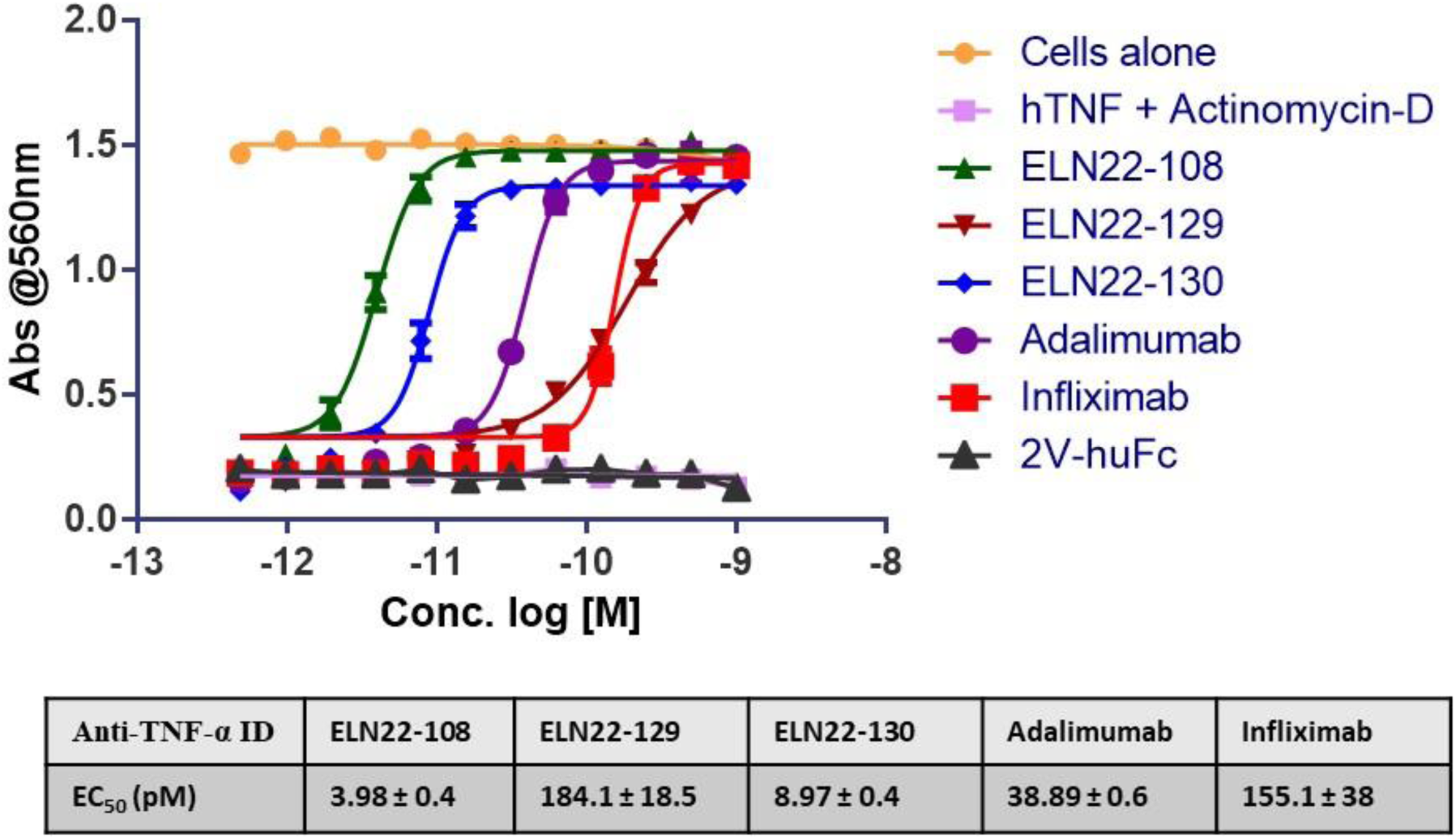
*In vitro* cell based potency characterisation of mono- and bi-paratopic Quad-X VNARs. Efficacy assessment in standard L929 cell-based assay. Neutralisation of 0.3 ng/mL hTNF-α-induced cytotoxicity in an actinomycin-D primed fibrosarcoma cell line (L929). Results are the mean ± SEM (n = 2, with 6 replicates). 2V-huFc is an isotype VNAR negative control.

**Table 1:** Summary of Octet R8 binding kinetics of anti-TNF-α mono- and bi-paratopic Quad-X constructs.

| Anti-TNF-α<br>ID | ELN22-108<br>(AHC2) | ELN22-108<br>(SAX) | ELN22-129<br>(AHC2) | ELN22-129<br>(SAX) | ELN22-130<br>(AHC2) | ELN22-130<br>(SAX) |
| --- | --- | --- | --- | --- | --- | --- |
| KD (pM) | 74.5 | 9.2 | 198 | 139 | 230 | 9.8 |
| K<sub>a</sub> (1/Ms) | $2.4 \times 10^5$ | $1.09 \times 10^6$ | $2.67 \times 10^5$ | $4.87 \times 10^5$ | $1.77 \times 10^5$ | $3.54 \times 10^5$ |
| K<sub>dis</sub> (1/s) | $1.7 \times 10^{-7}$ | $1 \times 10^{-5}$ | $5.3 \times 10^{-5}$ | $6.75 \times 10^{-5}$ | $4.07 \times 10^{-5}$ | $3.48 \times 10^{-6}$ |
| Full R<sup>2</sup> | 0.9884 | 0.9961 | 0.9989 | 0.9892 | 0.9993 | 0.9586 |

### Membrane TNF-α-mediated expression of E-selectin (CD62E) and IFN-γ in PHA activated CD4^+^ T cells

tmTNF-α mediated induction of the adhesion molecule, E-selectin (CD62E) in PHA activated CD4^+^ T cells was measured as a marker of reverse signalling. Unstimulated CD4^+^ T cells from healthy individuals did not express tmTNF-α, whereas treatment with 0.1 µg/mL PHA for 12 h resulted in the expression of tmTNF-α (data not shown). The treatment with acidified buffer to dissociate the cell-surface sTNF-α did not alter the intensity of the staining following binding of anti-TNF-α biologics to tmTNF. No E-selectin expression was detected in PHA stimulated or unstimulated conditions in the absence of a tmTNF-α binding mAb (data not shown), therefore confirming that E-selectin expression was dependent on an anti-TNF-α agent binding and activating tmTNF-α.^44,45^ MAB210 (R&D systems), a commercially available anti-human TNF-α binding and neutralising antibody, and Infliximab showed consistent and significantly superior surface tmTNF-α binding compared to the anti-TNF-α VNARs at 16 and 24 h timepoints (**Figure 6A-B**). The binding intensity of MAB210 and Infliximab significantly decreased at 48 h (**Figure 6C**) consistent with previously published data confirming internalisation of tmTNF-α bound anti-TNF-α antibodies.^46^ The observed surface binding kinetics also correlates with high levels of E-selectin induced at 24 h with both MAB210 and Infliximab (**Figure 6D**). Similar anti-TNF-α: tmTNF-α binding and E-selectin expression profiles were observed when the samples were analysed by flowcytometry (**Figures S3 and S4**). E-selectin was not expressed when anti-TNF-α test articles were immobilised on the assay plates prior to the addition of 0.1 µg/mL PHA stimulated CD4^+^ T cells (data not shown). The effect of these anti-TNF-α biologics on the induction of IFN-γ *via* tmTNF-α in PHA activated CD4^+^ T cells was then assessed. Previous reports have shown that IFN-γ is produced in response to an increase in monocyte apoptosis following tmTNF-α stimulation by anti-TNF-α antibodies.^40,47^ At the 24 h timepoint, MAB210 and Infliximab at 100 nM induced ∼ 10 – 15 ng/mL IFN-γ (Fig 6E, 6F). The anti-TNF-α VNARs were unable to induce any detectable levels of IFN-γ in these PHA stimulated CD4^+^ T cells (**Figure 6F**). VNAR-2V-huFc is an isotype control isolated from a naïve library and has no inherent binding property against any known target antigen.^48,49^

**Figure 6.**
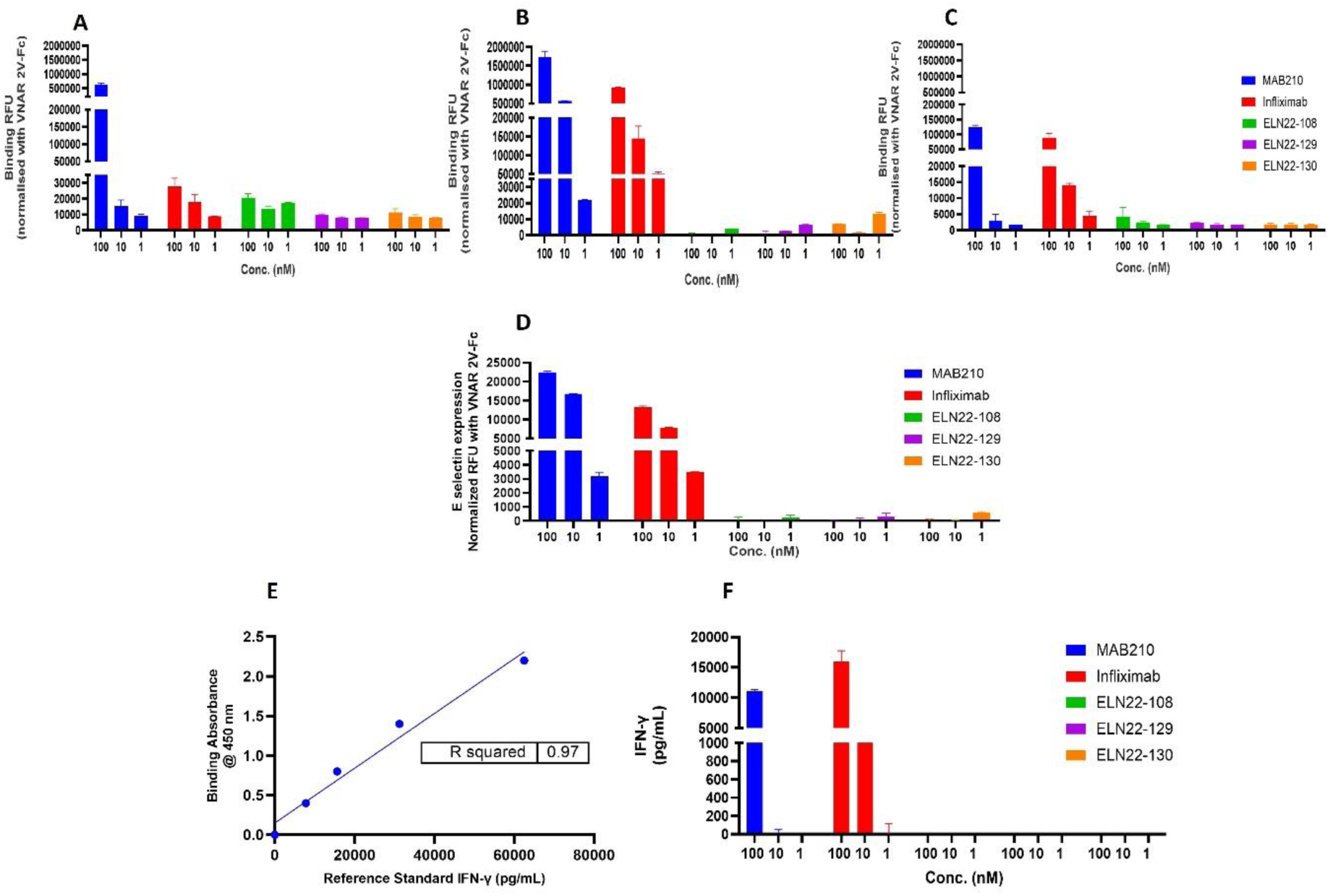
tmTNF-α dependent binding and downstream induction by anti-TNF-α agents on PHA activated normal human CD4^+^ T cells. (A, B, C) Fluorescence quantified surface binding of anti-TNF-α agents to tmTNF-α on PHA stimulated human CD4^+^ T cells at 16, 24 and 48 h timepoints respectively. The results are the mean ± SD (n = 2) with 2 replicates per experiment. Results were statistically analysed using a two-way ANOVA with Dunnett’s multiple comparisons *post hoc* test. At 24 and 48 h and 100 nM: Infliximab vs ELN22-108, ELN22-129 and ELN22-130 (*** *p < 0.0001*). MAB210 and Infliximab showed significant surface binding differences at 16 and 48 h with 100 nM dosing. (D) Fluorescence quantified E-selection expression at 24 h timepoint. The result is the mean ± SD (n = 2) with 2 replicates. Data were statistically analysed using a two-way ANOVA with Dunnett’s multiple comparisons *post hoc* test. At 24 h, Infliximab vs ELN22-108, ELN22-129 and ELN22-130 (** *p < 0.0035*). (E & F) IFN-γ secretion induced by the binding of anti-TNF-α agents to tmTNF-α on PHA stimulated CD4+ T cells were quantified using human IFN-γ DuoSet Sandwich ELISA kit. The results are the mean ± SD (n = 1) with 2 replicates per experiment, R^2^ = 0.97 for IFN-γ standard curve.

## Discussion

All five anti-TNF-α biologics (Adalimumab, Infliximab, Golimumab, Etanercept, and Certolizumab) currently in clinical use can neutralise TNF-α, however increasing evidence shows that all five drugs have differences in their therapeutic efficacy and utility and the occurrence of unwanted side effects. One possible explanation for at least some of these observed and well documented differences is how well they recognise tmTNF-α or more importantly the degree to which they can or cannot trigger full tmTNF-α reverse-signalling in tmTNF-α expressing cells.^8,10,11,50^ There is growing evidence implicating binding and induction of tm-TNF-α reverse signalling and associated tmTNF-α enabling mechanisms (i.e. CDC, ADCC) in the cytotoxic lysing and clearing of tmTNF-α bearing immune-surveillance T cells, antigen presenting cells, germinal centres, and memory B cells.^50–52^ These therapeutically unhelpful tmTNF-α mediated mechanisms can result in patients receiving treatment experiencing severe and life-threatening opportunistic infections and malignancies. For other patients their clinical history combined with the risk of these side effects, excludes them from ever receiving systemic anti-TNF-α therapies.

In this study, we presented two anti-TNF-α VNARs that recognized distinctly different epitopes on TNF-α. While VNAR-D1 bound a unique epitope not previously recognised by any of the existing anti-TNF-α agents, VNAR C4 recognises an epitope that overlaps with existing agents. VNAR-D1 and C4 have calculated contact areas of only 970 Å^2^ and 822 Å^2^ respectively (on the surface of each TNF-α protomer), which would be considered a small molecular footprint compared to the surface contact footprint of conventional antibodies on target proteins.^39,53,54^ Generally, larger surface area footprints result in increased complex stability, affinity, and neutralising potency. However, our mono-paratopic and bi-paratopic VNARs demonstrated equal or increased affinity and potency (*in vitro*) when compared to clinically approved anti-TNF-α biologics. This was achieved by packing multiple residue interactions with the CDR3 binding loops that protrude deep into their epitope on TNF-α.^55,56^

At physiologic concentrations, the TNF-α homotrimer slowly dissociates into monomers and reversibly trimerizes.^53,57–59^ Achieving robust neutralisation requires minimising the dissociation into monomers and stabilising the homotrimer state. Despite their small molecular footprint, VNAR-D1 and C4 effectively achieved stabilisation of the homotrimer. With the epitope of VNAR-D1 in the cleft of two protomers, three VNAR-D1 domains were able to bind simultaneously to the homotrimer (**Figure 2B**), Similarly, the VNAR-C4 epitope covered a single protomer that was accessible on all three protomers also allowing for binding in triplicate (**Figure 2A**). VNAR-D1 and C4 engagement with TNF-α stabilised the homotrimers by promoting interactions between the protomers and preventing a TNF-α monomer exchange reaction. This is like the protein:protein stabilisation achieved with Adalimumab and Infliximab.^39,53,56,60^ The low nanomolar binding affinities of 47 nM and 73 nM of VNAR-D1 and C4 monomers respectively correlates with the TNF-α homotrimer stabilisation. By the rational design of the bi-paratopic VNAR-D1 and C4, we improved this stabilisation resulting in several orders of magnitude enhancement in neutralising potency and binding affinity.^25^

The structural elucidation of VNAR-D1:TNF-α epitope implied a binding preference for sTNF-α over tmTNF-α. We were able to confirm both the absence of tmTNF-α binding *in vitro* and any induction of reverse signalling by activated CD4^+^ T cells. As markers of reverse signalling induction, we measured E-selectin and IFN-γ levels.^40,44,47^ There was no significant binding by any of the three anti-TNF-α VNAR formats tested (ELN22-108, ELN22-129 and ELN22-130), however, in stark contrast there was a time- and dose-dependent binding response seen with both MAB210 and Infliximab at 16 h, 24 h, and 48 h (Fig 6A, B, C respectively). Furthermore, the reduced binding signal measured for MAB210 and Infliximab at 48 h almost certainly correlates with the internalisation of these anti-TNF-α binding modalities. While the crystallography data did not suggest that VNAR-C4 may preferentially bind to sTNF-α, there was no measured binding to surface tmTNF-α on PHA activated CD4^+^ T cells at all timepoints. Possible explanations could be either rapid internalisation of the anti-TNF-α VNAR-C4 upon binding to tmTNF-α or because the smaller contact surface (epitope) on TNF-α recognised by this VNAR domain is conformationally different in both the transmembrane and soluble versions of TNF-α. The latter is more plausible since we do observe this level of flexibility in the structure of TNF-α. For instance the E-F loop of TNF-α plays a crucial role in the interaction with adalimumab and infliximab Fab fragments, however, this region is completely unobserved in the complex structures of TNF-α with TNFR2 or certolizumab, indicating that the E-F loop is flexible and not involved in these two interactions.^5,39,53,54^ An E-selectin expression time course has been reported to be different between CD4^+^ T cells induced by tmTNF-α with peak expression captured between 12 and 24 h.^44^ In this batch of CD4^+^ T cells, we were able to measure dose-dependent induction of E-selectin and IFN-γ expression above baseline at the 24 h timepoint for only MAB210 and Infliximab. Again, the mono-paratopic and bi-paratopic VNARs did not induce the expression of these molecules.

The bi-paratopic anti-TNF-α VNAR, ELN22-108 is capable of binding and neutralising sTNF-α *via* novel epitopes, with at least one of two epitopes believed to be an important differentiator between tmTNF-α and sTNF-α binding. In a previous study, we demonstrated that the bi-paratopic ELN22-108 is superior to Adalimumab in controlling sTNF-α-driven polyarthritis in Tg197 transgenic mouse model of polyarthritis, while in another study we showed evidence that VNARs targeting an unrelated antigen are amenable to binding unique epitopes that consequently resulted in novel mechanism(s) of antigen neutralisation.^24,31^ A combination of novel mechanism of action with enhanced safety profile and superior TNF-α neutralising potency positions ELN22-108 as an ideal first-line therapeutic for certain sub-sets of high-risk, autoimmune inflammation patients.

In summary, there is a strong case to be made for a novel class of anti-TNF-α targeting therapy with preference for sTNF-α or complete abrogation of tmTNF-α binding and induction. Also, in the absence of surface antigen binding, CDC and ADCC activation are impractical. It is understandable that such proposed mechanisms may have limited clinical efficacy benefit for patients with granulomatous diseases such as Crohn’s disease, although we believe there are additional approaches that could lead to optimal clinical outcomes for CD patients *via* sTNF-α-only targeting. This proposed mechanism will ensure enhanced safety profile for anti-TNF-α treated patients.

## Acknowledgements

This work was supported by the UKRI FLF/MRC Award MR/V026283/1 (O.U.), NIH/NCI R01 CA237272 (A.M.L.), NIH/NCI R01 CA233562 (A.M.L.), a 2018 Prostate Cancer Foundation Challenge Award (A.M.L.), a Prostate Cancer Foundation Young Investigator Award (A.M.L.), and Andy North and Friends (A.M.L.).

