## Supplemental Data for "The structural basis for the selective antagonism of soluble TNF-alpha by variable new antigen receptors"

**Table S1**

**Table S1: sTNF-α-VNAR-D1 data collection and refinement statistics**

| **Data collection** | TNF-α-VNAR-D1 |
|---|---|
| Source | P14, DESY |
| Space group | $P2_1$ |
| Cell (Å) | a = 112.32, b = 218.13, c = 236.49 |
| Angle (°) | α = 90.00, β = 96.28, γ = 90.00 |
| Wavelength (Å) | 0.9840 |
| Resolution limits (Å) | 109.30 – 3.31 (3.51 – 3.31) |
| No. of measured reflections | 547,405 |
| No. of unique reflections | 155,984 |
| $R_{merge}$ | 0.579 (2.725) |
| Mean [(I)/SD (I)] | 3.23 (0.57) |
| Multiplicity | 3.5 (1.9) |
| Completeness (%) | 92.8 (58.3) |
| Wilson B factor ($Å^{-2}$) | 48.25 |
| **Refinement** | |
| No. of monomers in the asymmetric unit | 60 |
| Resolution used in refinement (Å) | 109.30 – 3.31 |
| No. of unique reflections | 155,873 |
| Rfree (%) | 25.2 |
| Rwork (%) | 28.9 |
| No. of protein atoms (non-H) | 123500 |
| Protein atoms | 42.49 |
| RMSD bonds (Å)/angles (°) | 0.0228/2.9255 |
| Ramachandran favoured/outliers | 79.59/7.18 |
| Clash score | 28.63 |

Statistics for the highest-resolution shell are shown in parentheses.

**Table S2**

**Table S2: sTNF-α-VNAR-C4 data collection and refinement statistics**

| **Data collection** | TNF-□-VNAR-C4 |
|---|---|
| Space group | $C222_1$ |
| Unit cell dimensions | |
| *a, b, c* (Å) | 54.59, 89.45, 342.77 |
| Resolution (Å) | 46.6 - 1.92 (2.05 - 1.92) |
| $R_{sym}$ or $R_{merge}$ | 0.099 (0.883) |
| $I / \sigma I$ | 8.8 (1.8) |
| Completeness (%) | 89.3 (62.0) |
| Redundancy | 5.3 (5.0) |
| $CC_{1/2}$ | 0.997 (0.563) |
| **Refinement** | |
| Resolution (Å) | 46.6 - 1.92 (1.99 - 1.92) |
| No. Reflections | 48613 (706) |
| $R_{work}$ / $R_{free}$ | 0.198/0.246 (0.246/0.349) |
| No. atoms | 6397 |
| Protein | 6143 |
| Ligand/ion | 50 |
| Water | 204 |
| *B*-factor | 49.67 |
| Protein | 49.62 |
| Ligand/ion | 71.50 |
| Water | 45.79 |
| Ramachandran plot | |
| Favored (%) | 98.56 |
| Allowed (%) | 1.44 |
| Outliers (%) | 0.00 |
| R.m.s. deviations | |
| Bond lengths (Å) | 0.016 |
| Bond angles (º) | 1.07 |

Statistics for the highest-resolution shell are shown in parentheses.

**Supplementary Fig 1**

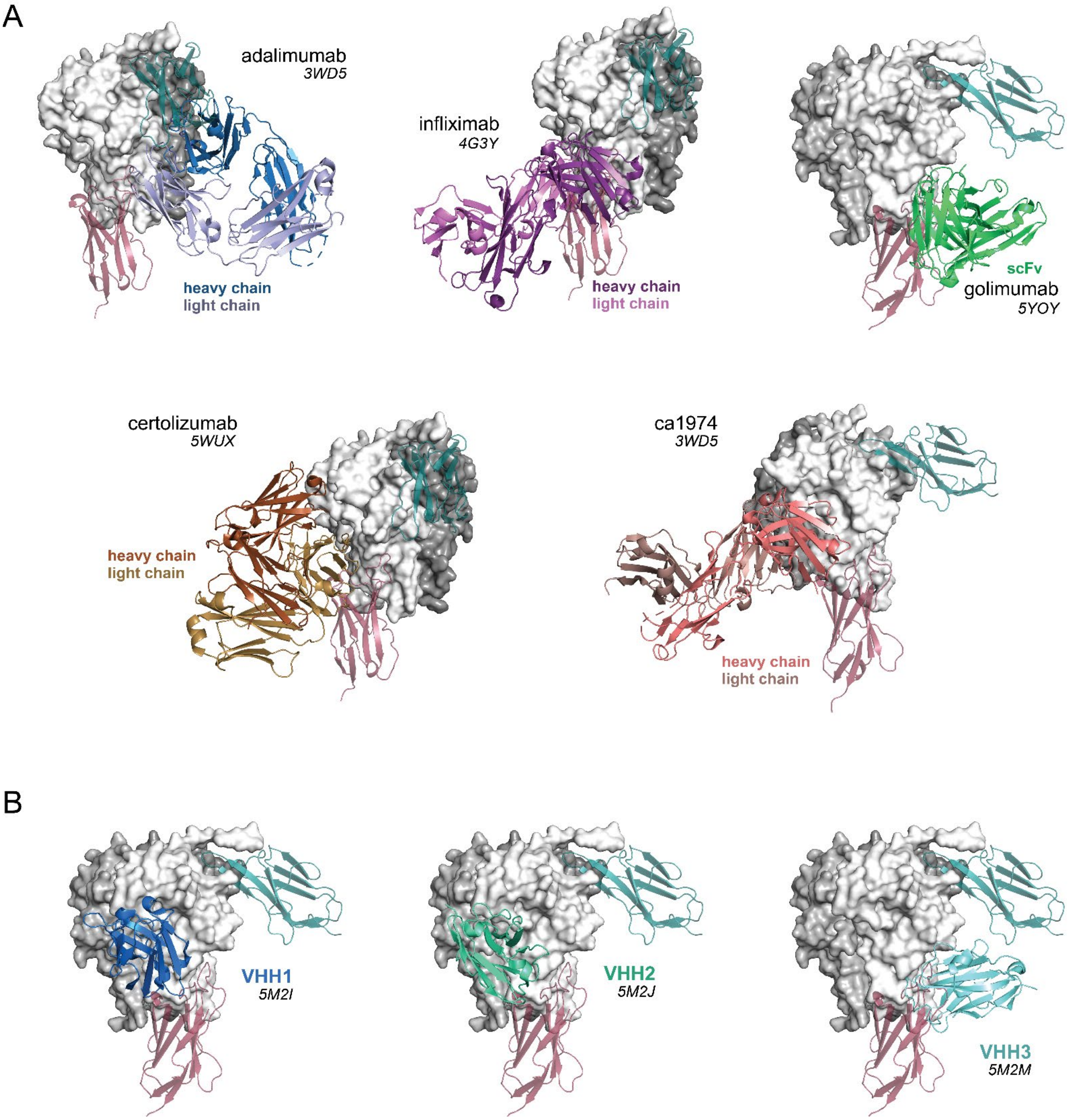


**Figure S1. Comparison of VNARs and existing biological epitopes.** A, B) The figure shows a grayscale surface representation of the TNF-α trimer bound by a monomer of VNAR-C4 (red) and VNAR-D1 (green). An epitope comparison is presented by overlaying the crystal structures of available TNF-α biologics. Multi-chain structures are show in Panel A, and nanobody structures are shown in Panel B.

**Supplementary Figure 2**

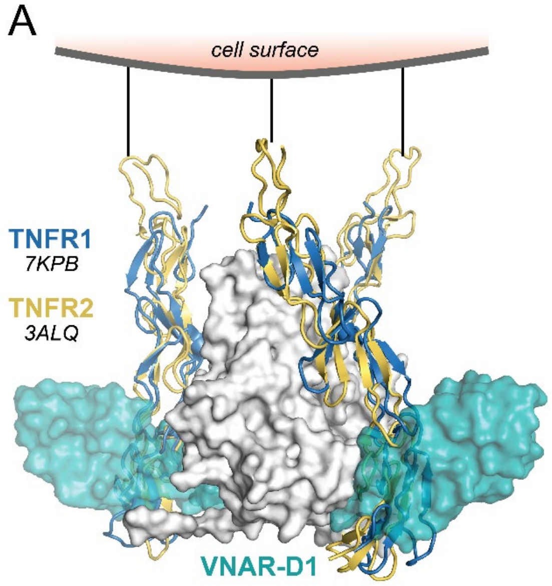


**Figure S2. VNAR-D1 blocks TNFR binding.** A) Overlaid crystal structures of TNFR1 (blue) and TNFR2 (yellow) shown as cartoons bound to a surface representation of the TNF- α trimer (grayscale) with an illustration indicating the receptor position relative to the cell surace, as in Figure 3A. A surface representation of VNAR-D1 has been superimposed onto the structures showing the overlap with the TNFRs.

### Supplementary Figure 3

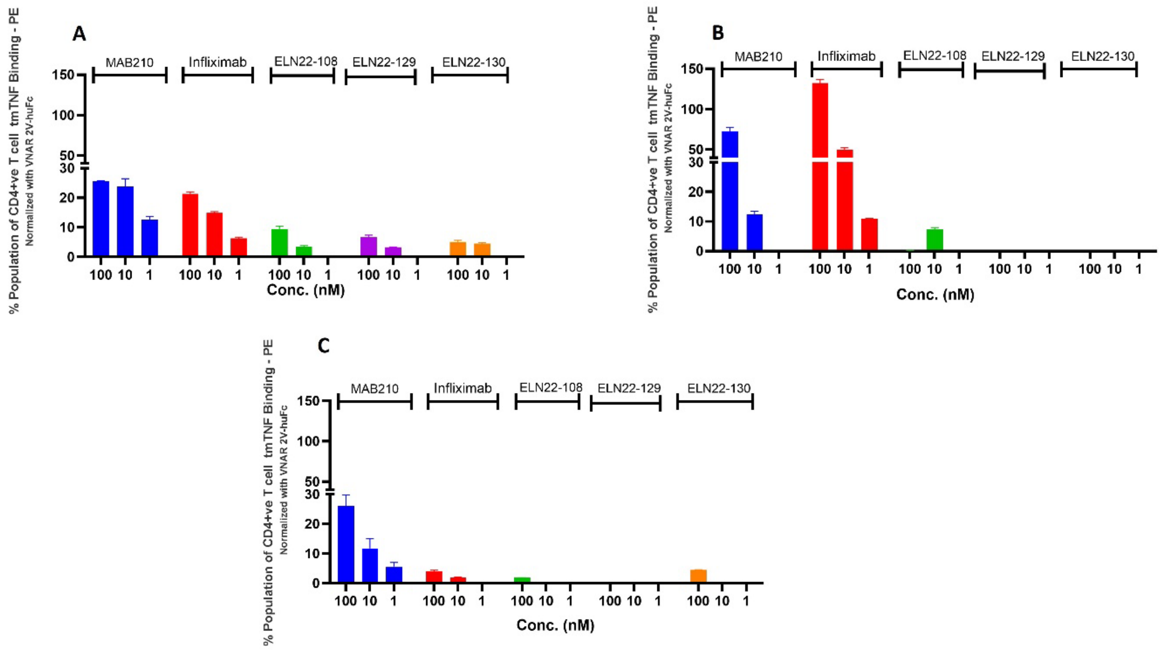


**Figure S3. tmTNF-α dependent binding of anti-TNF-α agents on PHA activated normal human $CD4^+$ T cells.** (A, B, C) Flowcytometry quantified surface binding of anti-TNF-α agents to induced tmTNF-α on PHA stimulated human $CD4^+$ T cells at 16, 24 and 48 h timepoints respectively. The results are the mean ± SD (n = 2) with 2 replicates per experiment. Results were statistically analysed using a two-way ANOVA with Dunnett's multiple comparisons *post hoc* test. At 16 and 24 h, Infliximab at 1, 10 and 100 nM vs ELN22-108, ELN22-129 and ELN22-130 (*** $p < 0.0001$).

### Supplementary Figure 4

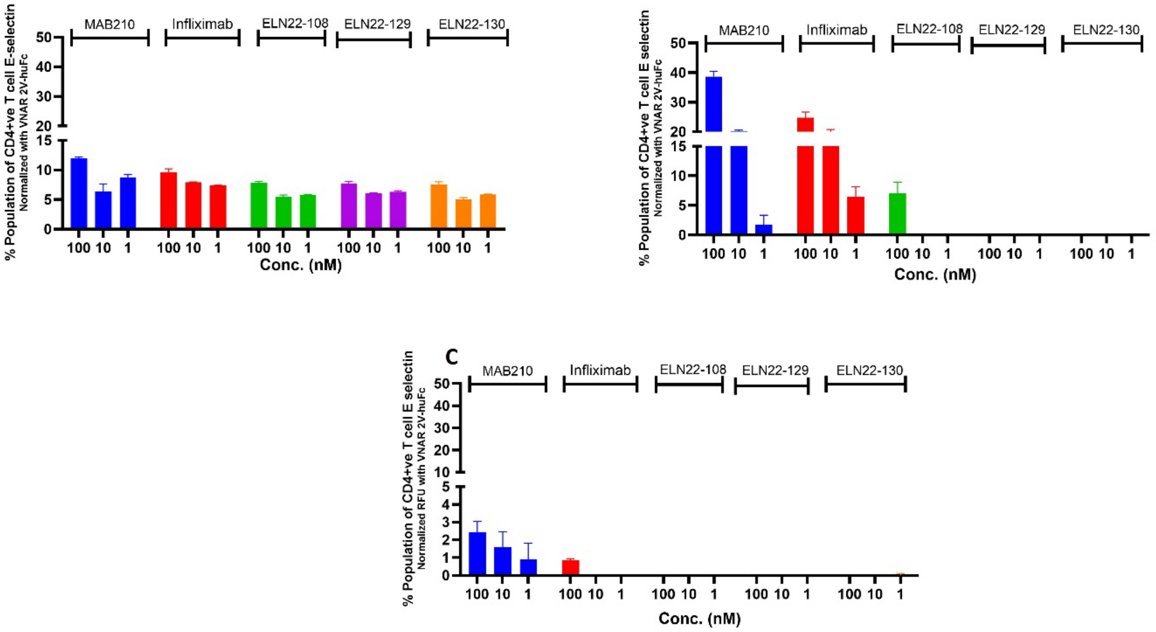


**Figure S4. Anti-TNF-α induction of E-selectin via tmTNF-α mediated "outside-to-inside" signalling in PHA activated normal human CD4$^+$ T cells.** (A, B, C) Flowcytometry quantified E-selection expression at 16, 24 and 48 h timepoints respectively. The results are the mean ± SD (n = 2) with 2 replicates per experiment. Results were statistically analysed using a two-way ANOVA with Dunnett's multiple comparisons *post hoc* test. At 16 and 24 h, Infliximab at 1, 10 and 100 nM vs ELN22-108, ELN22-129 and ELN22-130 ($p < 0.0018$). At 48 h and 100 nM, Infliximab vs ELN22-108, ELN22-129 and ELN22-130 (* $p < 0.0338$).
